# Centimeter-scale, physiologically relevant intestinal organoids generated entirely from pluripotent stem cells

**DOI:** 10.1101/2024.08.15.608057

**Authors:** Zhen Qi, Zhanguang Zuo, Yi Dong, Jingyu Shao, Chong Wang, Rosanna Zhang

## Abstract

Over the past decade, intestinal organoids have shown great promise as a platform to study the development of intestine, elucidate the pathogenesis of diseases such as inflammatory bowel disease (IBD), and model physiological features of intestinal tissue for high-throughput drug screening. However, intestinal organoids derived from adult epithelial stem cells lack cellular diversity, especially those resides in the lamina propria. Intestinal organoids derived from embryonic stem (ES) cells or induced-pluripotent stem cells (iPSCs) has greater cellular diversity, but are still limited in size and physiological features such as peristalsis. In this study, we generated centimeter-scale, full-thickness, and physiologically relevant intestinal organoids in suspension culture without usage of Matrigel. Using a series of optimized culture media, multiple lineages of cells were differentiated from iPSCs and spontaneously assembled to achieve the intestinal architecture. These bubble-like organoids have a thickness of 500 microns, exhibit a mature vasculature network, and have smooth muscle-like cells to conduct regular peristalsis. In addition, adipocyte-like cells and granulocyte-like cells are also observed in these organoids, which are important in immune homeostasis. Lastly, these organoids show mature crypt structures, response to lipopolysaccharides (LPS), and increases luminal influx upon forskolin treatment, suggesting the organoids have intact epithelial integrity. Thus, this study provides a highly reproducible approach to produce large and physiologically relevant intestinal organoids that are suitable for different biomedical applications.

## Introduction

Intestinal organoids are three-dimensional (3D) in vitro cell cultures that closely mimic the physiological structure, biological complexity, and key functions of the human intestine (Sato, Stange et al. 2011, Clevers 2016, Fatehullah, Tan et al. 2016). These organoids have become a valuable tool for studying intestinal development and pathology (Huch, Knoblich et al. 2017). In addition, they offer a more accurate platform for drug development compared to traditional 2D cell cultures, enabling high-throughput screening and reducing the reliance on animal models (van de Wetering, Francies et al. 2015, Vlachogiannis, Hedayat et al. 2018, Schutgens and Clevers 2020). Intestinal organoids support advancements in personalized medicine, gene and cell therapies, and regenerative medicine (Lancaster and Knoblich 2014, Bredenoord, Clevers et al. 2017, Kim, Koo et al. 2020, Liu, Li et al. 2024), suggesting great potentials for different biomedical needs.

A typical intestinal organoid consists of a single layer of polarized intestinal epithelial cells surrounding a central lumen, with crypt-villus structure located on the lumen side (Sato, Vries et al. 2009). Intestinal organoids with such features have been used to study intestinal nutrient transport, drug absorption and delivery, incretin hormone secretion, and infection by various enteropathogens (Finkbeiner, Zeng et al. 2012, Hill and Spence 2017). Intestinal organoids can be generated directly from intestinal stem cells, intestinal tissues or pluripotent stem cells in vitro and maintained in a serum-free medium supplemented with essential factors such as EGF, the Wnt agonist R-spondin, and the BMP inhibitor Noggin (EWRN) (Spence, Mayhew et al. 2010, Sato, Stange et al. 2011, Yui, Nakamura et al. 2012). Despite of self-maintained proliferation and differentiation of epithelial lineages, these organoids must be maintained in Matrigel and lack essential lamina propria components. To resolve this, multi-lineage intestinal organoids were generated using adult intestinal tissue or iPSC differentiated cells, including a variety types of cells such as mesenchymal cells, enteric neuronal cells, vascular cells, and immune cells can be provided (Spence, Mayhew et al. 2010, Watson, Mahe et al. 2014). However, these intestinal organoids are normally less than 1 mm, and hence they cannot be used to reflect the dimension of real intestinal organ tissue. In addition, lacking peristalsis capability in existing intestinal organoids results in only passive diffusion-derived substance exchange between the lumen side and the outer aqueous environment. An improved method is therefore needed to generate large intestinal organoids with physiologically relevant functions to better recapitulate the intestine *in vivo*.

In this study, we developed a cell culture kit, which is optimized to differentiate iPSCs into intestinal organoids with multiple cellular lineages and peristalsis capability. With continuous culture, the size of the intestinal organoids increased over time and achieved a centimeter-scale upon 60 days. These organoids displayed a full-thickness architecture with a typical central lumen and crypt-villus structure. In addition, these organoids also exhibited typical physiological functions such as fluid efflux. Moreover, immune responses were evoked by the addition of LPS in the culture medium, leading to observable organoid atrophy. This study verified the feasibility of using a simple kit to develop iPSCs into mature intestinal organoids with high physiological relevance, high reproducibility, and multiple functions.

## Methods

### 1 Generating iPSC-derived intestinal organoids

iPSCs (ATCC, ACS-1007) were cultured under mTeSR (STEMCELL Technologies, 100-0276) culture system in 6-well TC-treated Plates (Corning, 3516) coated with 1% Matrigel (Corning, 354277). When the cell confluency is greater than 90%, iPSCs can be readily used for intestinal organoid induction. The intestinal organoid differentiation is conducted by using Human iPSC-Derived Intestinal Organoid Differentiation Kit (ACROBiosystems, RIPO-IWM005K-1kit) and following the instruction. Simply, iPSCs are switched to pre-warmed intestinal medium A and the medium is changed every day for 3 days (total 72 h). Then, change medium to intestinal medium B for 9 days (change medium every other day). On day 12, cells are detached using TrypLE (Gibco, 12604013) and adjusted to 2.5×10 cells/mL in Intestinal Medium C. The 200 μL of cells (approximately 50,000 cells/well) are then settled in an ultra-low attachment U-shape 96-well plate (Corning, 7007) for 72 h to allow spheroid formation. Lastly, spheroids were transferred into ultra-low attachment 6-well plates (Corning, 3471) containing intestinal medium D for expansion. At day 45 from differentiation, change the medium into Intestinal organoid maturation medium (ACROBiosystems, RIPO-IWM006) (medium change every other day). Intestinal organoids will start to peristalsis from day 60 of differentiation and reach 100% of peristalsis at day 80.

### 2 Immunofluorescence staining of intestinal organoids

Organoids were fixed in 4% PFA, embedded in paraffin, and cut into 4 to 6 μm sections, followed by deparaffinization and heat-mediated antigen retrieval treatment (Histova Biotechnology, AbCracker 2in1 Retrieval/Fast Elution Solution, ABCFR5L). Multiplexed immunofluorescence assay was conducted as previously described (Ye, Zhou et al. 2021). Multiplexed IF slides were imaged using a confocal laser scanning microscope (Zeiss, LSM880).

### 3 Evaluating the epithelial functions of intestinal organoids

Forskolin treatment: 100-day intestinal organoids were treated with 1 μM, 5 μM, 10 μM forskolin for 6 days in Advanced DMEM/F-12 (Gibco, 12634010)(Medium change every other day). Photos were taken on days 0, 2, 4, and 6, and organoid diameters were measured using ImageJ. LPS treatment: 50-day intestinal organoids were treated with 1 μg/ml and 10 μg/ml LPS for 18 days in Advanced DMEM/F-12. After LPS treatment, organoids were collected and lysed for total RNA extraction and RT-qPCR.

### 4 Absorption of fatty acid in intestinal organoids

34-day intestinal organoids were stained with 5 μM Dil (Yeasen, DilC18(3)) in Advanced DMEM/F-12 for 10-15 min at 37 °C, 5% CO_2_. After staining, organoids were washed with DMEM for 3 times (2 min each time). Then organoids were treated with 5 μM BODIPY 500/510 (ThermoFisher, C1-BODIPY-C12) for 2 h (at 5% CO_2_, 37 °C). After incubation, organoids were washed and fixed with 4% PFA overnight. Fluorescent images were captured using an inverse fluorescence microscope (Discover Echo, Revolve Generation 2).

### 5 Bulk RNA-seq of intestinal organoids

Intestinal organoids were collected and lysed for total RNA extraction using Trizol reagent (Invitrogen, 15596026). RNA quality was confirmed using a NanoDrop and a Bioanalyzer (Agilent, 2100). RNA libraries were constructed using the TIANSeq Fast RNA Library Prep Kit (Illumina) (TIANGEN Biotech, NR102) and sequenced using PE150 settings. Gene expression levels were then analyzed using HTSeq and reported in FPKM. Differential gene expression (DGE) analysis was conducted using edgeR (version 3.32.1, R version 4.0.3) on the read count data.

### 6 Statistical analysis

Statistical significance was assessed using one-way ANOVA, followed by post-hoc test with Dunnett’s test for multiple comparisons against control group or Tukey’s test for multiple comparisons between all pairs of groups. Significance is indicated by asterisks: * for p < 0.05, ** for p < 0.01, and *** for p < 0.001.

## Results

We first optimized cell culture procedure to differentiated iPSCs into intestinal organoids using our commercialized cell culture media (Fig.1A). To begin with, iPSCs were cultured as adherent cells in medium A for 3 days followed by 9 days in medium B. Afterwards, cells were dissociated and cultured in medium C as suspension cells (50,000 cells/ 200 ul) for 3 days to allow organoid formation. Spherical organoids were further differentiated in medium D for 30 days. At this time point, these organoids exhibited a more developed state with internal cavity structure. Organoids can then be cultured in maturation medium for up to 40 days. By day 64, the diameter of these bubble-like intestinal organoids can reach 5 mm on average (Fig.1C and D).

**Figure 1.**
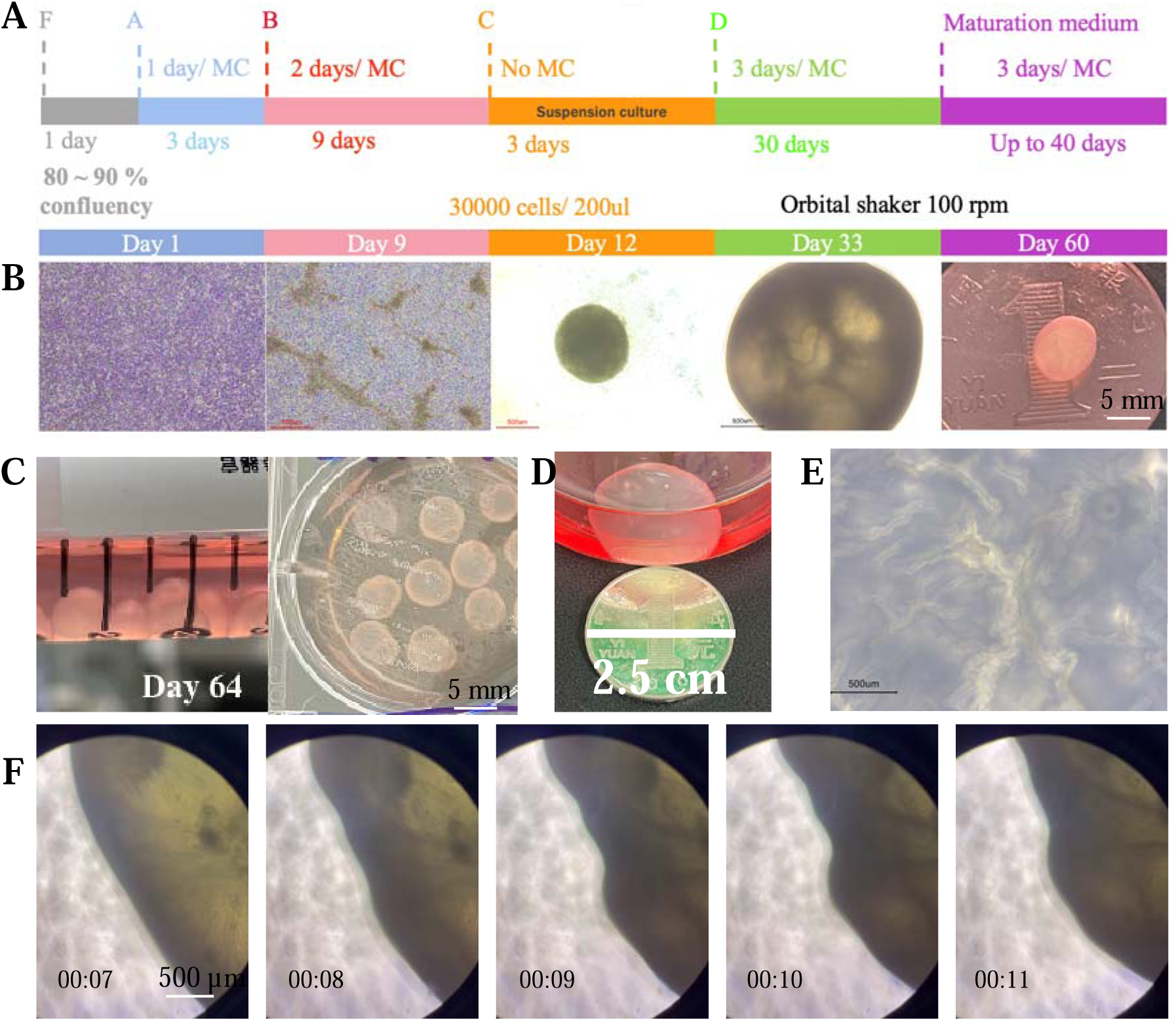
Induction and Maturation of Intestinal Organoids from iPSCs. (A) Schematic plot showing the culture procedure used to form intestinal organoids from iPSCs. (B-C) Representative images of intestinal organoids at different differentiation stages. (D) Forskolin-treated (forskolin 10 μM, 5 days) human intestinal organoids. (E) The fold structure of intestinal organoids on the lumen side after being cut open. (F) Representative images showing the rhythmic peristalsis of organoids. Time were shown in min : sec.

In addition to rapid differentiation, we also observed that the developed organoids have rhythmic peristalsis (Fig.1F, supplemental movie S1 and S2), which is similar to the rhythm observed in human intestine (Huizinga, McKay et al. 2006). This result suggests that our intestinal organoids can simulate basic movement of native intestine organ, providing additional features that were not reported in other intestinal organoid systems (Sato, Vries et al. 2009, Spence, Mayhew et al. 2011, Workman, Mahe et al. 2017).

During embryonic development, the digestive track initiates from week 3-4 and divides into three distinct sections: foregut, midgut, and hindgut (Bhatia, Shatanof et al. 2024). To better characterize the developmental stage and the part of the intestine that we generated, we dissected a 120-day organoid (Fig.1E and supplemental movie S3.) and performed histological analysis (Fig.2). We first observed a large empty cavity in the luminal side of the organoid, which is similar to the lumen of the intestine. We then observed polarized epithelial cells with crypt structures and microvilli on some epithelial cells (blue dotted region), a feature that is commonly observed in the small intestine (Fig. 2C).

**Figure 2.**
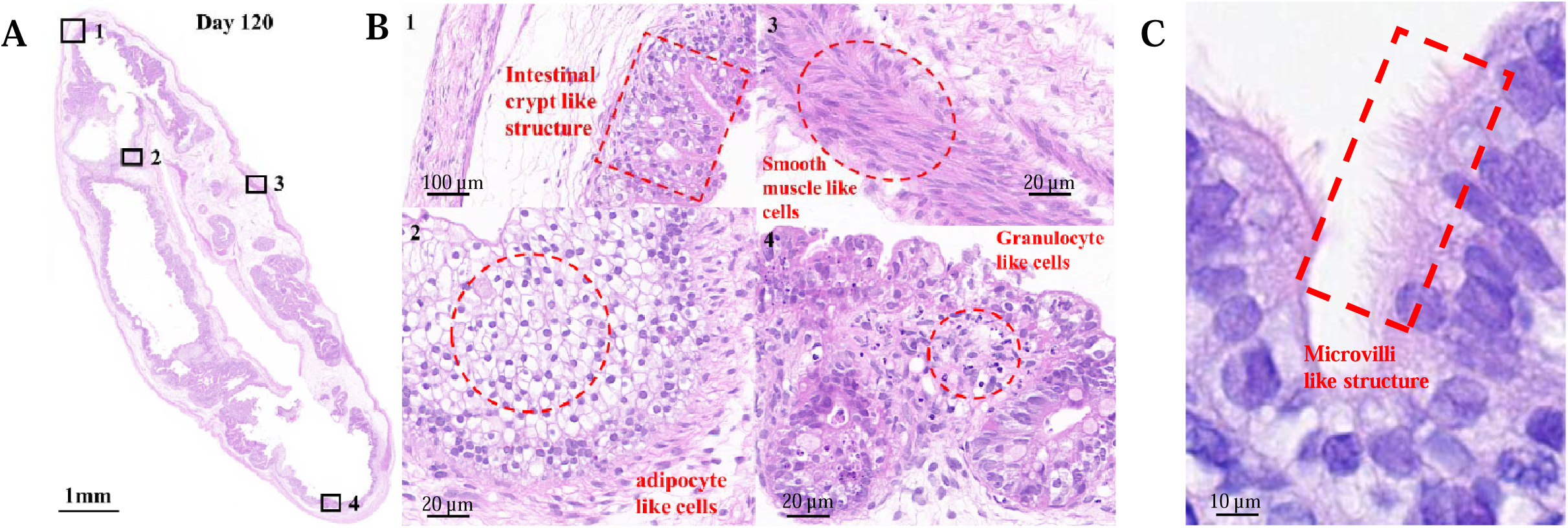
Structural Features of Matured Intestinal Organoids. (A) Cross-sectional structure shown by H&E staining of intestinal organoids. (B) Detailed view of local structures after HE staining of intestinal organoids. (C) Detailed view of microvilli-like structures.

Next, we investigated the stromal part of the organoid. We observed several cell types in the tissue, including fibroblasts, smooth muscle-like cells, granulocyte-like cells, and adipocyte-like cells. Among these, fibroblasts play important roles in maintaining tissue architecture and supporting different cell types. Smooth muscle-like cells are important for bowel movement, which may produce rhythmic peristalsis that we observed. Granulocyte-like cells may recapitulate tissue resident macrophages that play an important role in innate immune responses. Taken together, these cell types suggest a successful differentiation of iPSCs into various cell types typical of the intestinal tissue.

To further characterize the cell types observed in these intestinal organoids, we performed immunofluorescence staining and observed several subsets of epithelial cells, including MUC2^+^ goblet cells (Fig. 3A, 3C), CHGA^+^ enteroendocrine cells (Fig. 3C), and Ki67^+^ proliferating cells (stem cells and transit amplifying cells, Fig. 3C). In addition, we observed α-SMA^+^ stromal cells underneath the epithelial layer (Fig. 3B) and CD68 macrophage-like cells in the lamina propria (Fig. 3B, 3C). Surprisingly, we also observed CD31^+^ endothelial-like cells that formed vascular-like structures and networks in the stromal layer (Fig. 3D, 3E). These observations further suggest that iPSCs were differentiated into many important cell types and these iPSC-derived intestinal organoids can recapitulate many aspects of the intestine.

**Figure 3.**
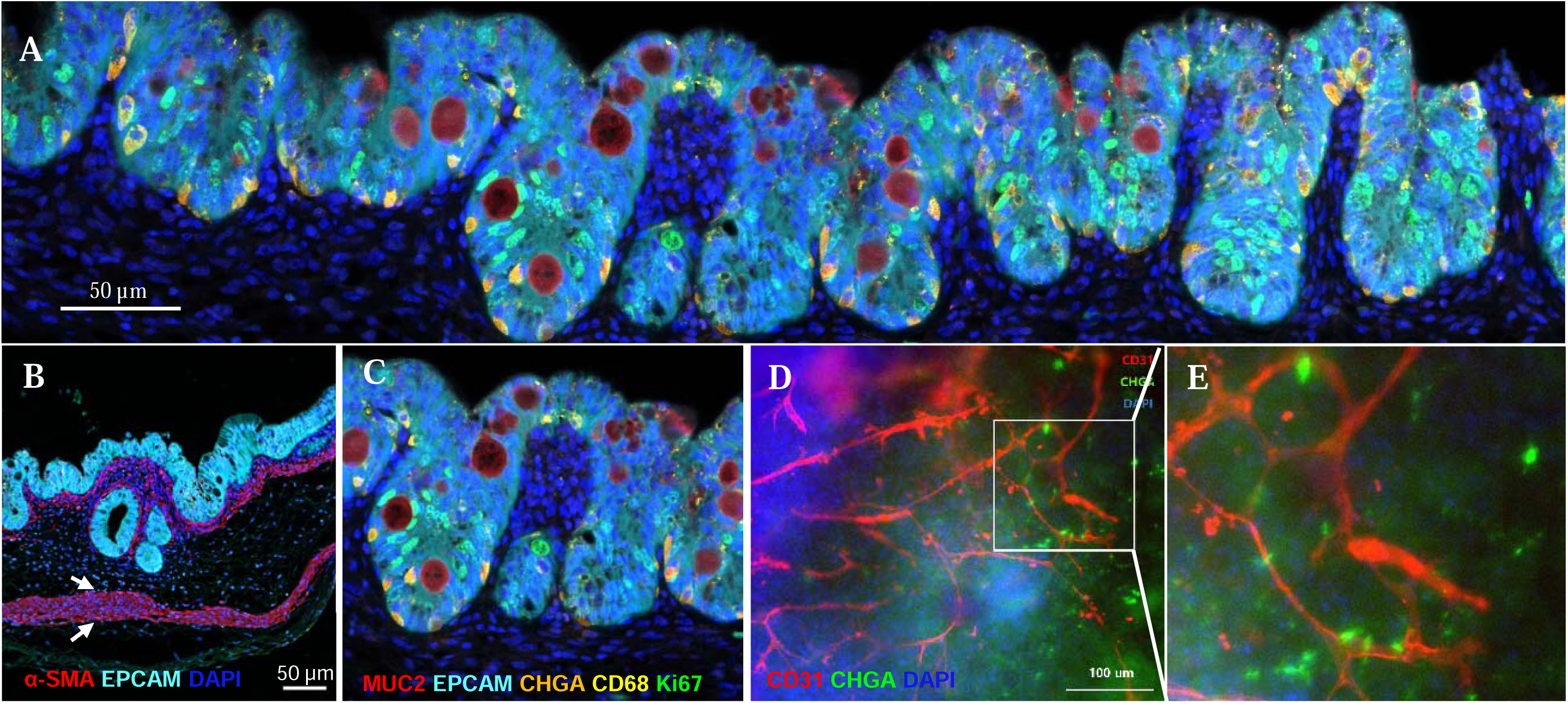
Immunofluorescence Staining of Intestinal Organoids. (A) Cross-section view of an intestinal organoid. (B-E) Detailed view of local structures and cell types after immunofluorescence staining of intestinal organoids. EPCAM, epithelial cell marker; MUC2, goblet cell marker; CHGA, enteroendocrine cell marker; Ki67, proliferating cell marker; α-SMA, myofibroblasts and smooth muscle cell marker; CD68, macrophage cell marker; CD31, endothelial cell marker.

Next, we compared the cellular diversity of organoids on differentiation day 30 and day 61. Using RNA sequencing, we found 2041 upregulated and 1758 downregulated genes (Fig. 4A). Among these, we observed an overall increasing trend of epithelial cell diversity during maturation (Fig. 4B). The transcriptional expression levels of the markers of Enterocytes (VIL and SOX9), Paneth cells (LYZ and MMP7), goblet cells (MUC2), and stem cells (LGR5) increased on day 61 compared to day 30. These results were consistent with our observation that intestinal organoids gradually increased their cellular diversity in the maturation media.

**Figure 4.**
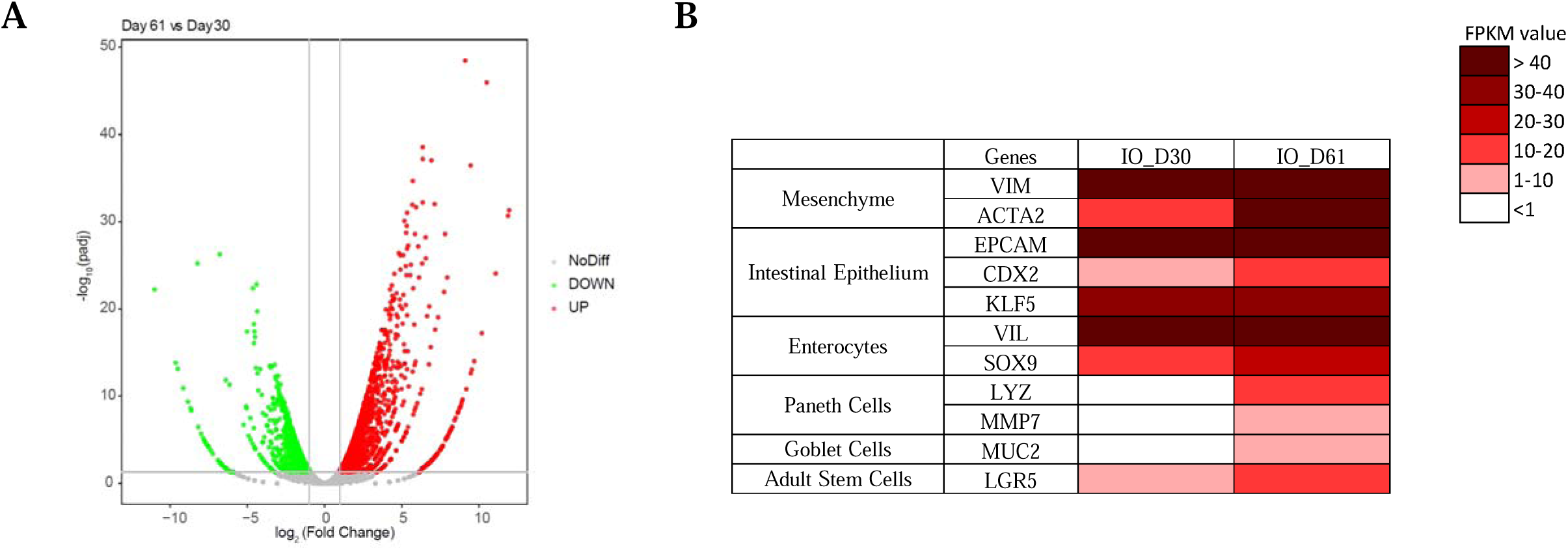
Gene Expression Analysis of Intestinal Organoids. (A) Differential gene expression analysis between 30- and 61-day organoids. (B) RNA-seq analysis comparing the expression levels of genes representing different subsets of intestinal cell types between 30- and 61-day organoids.

As an important digestive organ, the small intestine is responsible for absorbing nutrients, including fatty acids (Basile, Launico et al. 2024). We first conducted a fatty acid diffusion assay to valid the biological function of our intestinal organoids. We added C1-BIDIPY^+^ fatty acid to the culture media and observed that the fatty acid can penetrate the cell membrane and diffuse to the luminal side of the organoid. This result suggest that our intestinal organoids have basic functions in fatty acid transport and may be used for other absorption assays.

Other important features of the intestine is the barrier function and fluid transport capability (Groschwitz and Hogan 2009). We next treated organoids with forskolin, an activator of cystic fibrosis transmembrane conductance regulator (CFTR), an important chloride channel (Boj, Vonk et al. 2017). By treating organoids with forskolin at different concentrations, we observed a dose-dependent increase in the organoid volume (p<0.05), due to osmolarity balance (Fig. 5B). This result suggests that our intestinal organoids have intact barrier function and can transport fluid upon ion channel activation.

**Figure 5.**
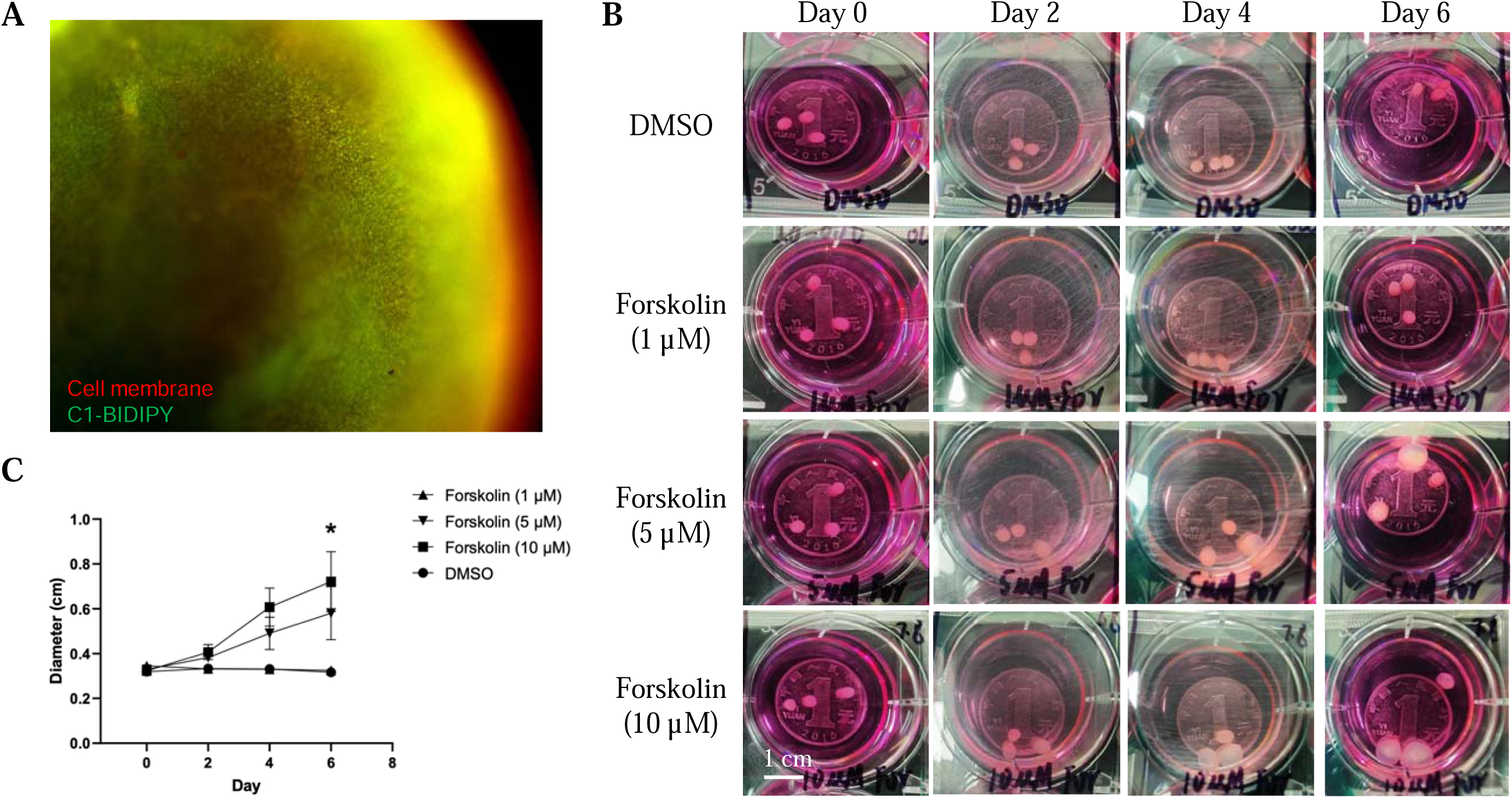
Absorption and barrier functions of intestinal organoids. (A) R representative image showing fatty acids absorption by intestinal organoids. (B) Representative images showing forskolin-induced swelling of intestinal organoids. (C) Quantitative changes in diameter of intestinal organoids. One data point represents the mean of n = 3 organoids. Statistical significance was assessed using one-way ANOVA on day 6, followed by post-hoc test with Dunnett’s test for multiple comparisons. Asterisks indicate significant differences between forskolin 10 μM and DMSO group (p < 0.05).

Since barrier disruption and bacteria invasion can trigger many intestinal diseases (Ohland and Jobin 2015), we then examined the immune responses of our organoids when encountered LPS. A volume reduction of the organoids was first observed (Fig. 6A), which could be likely attributed to the leakage of luminal fluids (Guo, Al-Sadi et al. 2013). By measuring the cytokine secretion of the organoids, a significant and dose-dependent increase of pro-inflammatory cytokines IL-6 and CCL2 (Fig. 6B) was detected (p<0.05). Such two cytokines are commonly observed during bacterial infection in the intestine (Yagi, Takaki et al. 2002, Sonnier, Bailey et al. 2012). All these results demonstrating the functional relevance of our intestinal organoids in modeling intestinal infection and inflammation.

**Figure 6.**
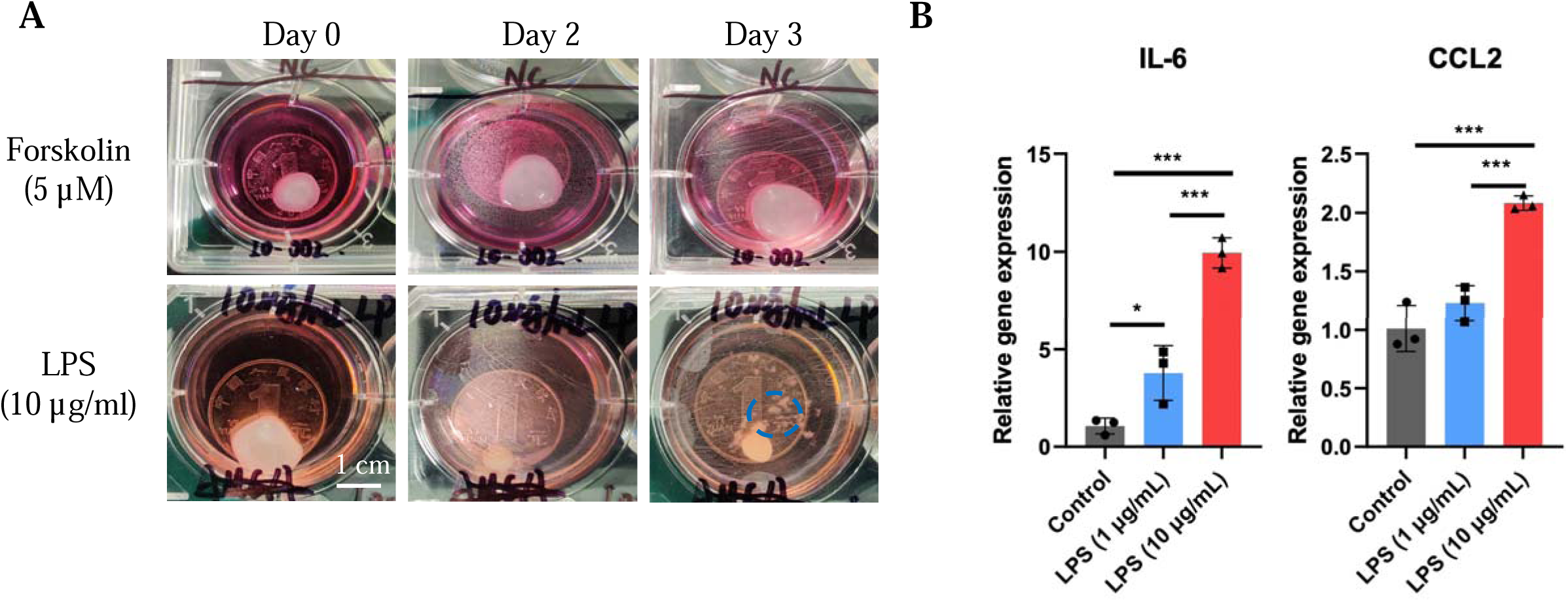
Immune responses of intestinal organoids. (A) Representative images showing a decrease in the size of organoids after LPS treatment. (B) Cytokine responses of organoids after LPS stimulation. Each data point represents one organoids, n = 3. Post-hoc analysis used Tukey’s multiple comparisons test, * p < 0.05, ** p < 0.01, *** p < 0.001.

## Discussion

In this study, we show a robust organoid model that recapitulates many aspects of the natural intestine organ *in vivo*. Comparing with other studies that used either adult epithelial stem cells or iPSCs, our method have two major benefits. First, our intestinal organoids are derived from iPSCs that is highly reproducible and scalable, making it possible for high throughput studies. Second, our intestinal organoids recapitulate more physiologically relevant features than previous reported organoids (Sato, Vries et al. 2009, Spence, Mayhew et al. 2011, Workman, Mahe et al. 2017), including greater organoid size, presence of endothelial-like cells, macrophage-like cells, and more importantly, spontaneous rhythmic peristalsis. Childs et al. observed peristalsis in human intestinal organoids that were first cultured in a murine host for 30 days (Childs, Poling et al. 2024). However, the rhythm in their organoids was much slower (2 beats per minute) than ours (3-4 beats per minute) and host stimulation was required. Although we did not focus on characterizing neuronal cells in our intestinal organoid, as reported previously (Workman, Mahe et al. 2017), we believe our intestinal organoid also includes functional neurons because rhythmic peristalsis requires neuronal regulation of muscle cells (Spencer and Hu 2020). All these observations suggest that our intestinal organoids can recapitulate the complex physiological features of the natural intestine organ.

Since our intestinal organoid recapitulates many cell types and features of the natural intestine, it can be used as an advanced platform for disease modeling. A typical example of this would be inflammatory bowel disease (IBD), especially Crohn’s disease (CD). IBD is a multifactorial disease with complexed etiology (Uhlig and Powrie 2018), which requires the involvement of multiple cell types (Chang 2020, Dong, Johnson et al. 2024). For CD, in additional to disruption of the epithelial cells, fibrosis is also an important symptom that disrupts the mesenchymal layer (Uhlig and Powrie 2018), which cannot be recapitulated by epithelial-only organoids. Our intestinal organoids have both mucosa and submucosa. It can therefore be an excellent pathological model to study CD and many other intestinal diseases.

In addition to the biological advantages of our intestinal organoids, we also simplified the procedure of generating organoids by pre-formulating the differentiation media. Previous studies use freshly made media that requires reconstitution of different growth factors and chemicals (Spence, Mayhew et al. 2011, Workman, Mahe et al. 2017), therefore the chance of batch variations is high due to handling issues and materials from different vendors. Our media is commercialized, hence we can reduce a lot of factors that leads to batch variations. This will also improve the reproducibility of organoid studies.

## Conclusion

In this study, we successfully developed centimeter-scale, mature, physiologically relevant intestinal organoids from iPSCs using our commercialized media products. These organoids exhibited structural complexity, periodic peristalsis, and functional capabilities, making them a valuable model for studying intestinal physiology and diseases. Our intestinal organoids recapitulated more features of the intestine than existing organoid models (Sato, Vries et al. 2009, Spence, Mayhew et al. 2011, Workman, Mahe et al. 2017), making it a more accurate representation of native intestinal tissue.

## Acknowledgments

We would like to thank Mike Chen for providing the fund for this research.

## Author Contributions

Conceptualization: Z.Q.;

Experimental investigation: Z.Q., Y.D. and Z.Z.;

data analysis: Z.Q., Z.Z., Y.D. and J.S.;

original draft: C.W. and Y.D.;

revision: Y.D., Q.Z., Z.Z., C.W. and R.Z.

## References

Basile, E. J., M. V. Launico and A. J. Sheer (2024). Physiology, Nutrient Absorption. StatPearls. Treasure Island (FL), StatPearls Publishing Copyright © 2024, StatPearls Publishing LLC.

Bhatia, A., R. A. Shatanof and B. Bordoni (2024). Embryology, Gastrointestinal. StatPearls. Treasure Island (FL), StatPearls Publishing Copyright © 2024, StatPearls Publishing LLC.

Boj, S. F., A. M. Vonk, M. Statia, J. Su, R. R. Vries, J. M. Beekman and H. Clevers (2017). “Forskolin-induced Swelling in Intestinal Organoids: An In Vitro Assay for Assessing Drug Response in Cystic Fibrosis Patients.” J Vis Exp(120).

Bredenoord, A. L., H. Clevers and J. A. Knoblich (2017). “Human tissues in a dish: The research and ethical implications of organoid technology.” Science 355(6322).

Chang, J. T. (2020). “Pathophysiology of Inflammatory Bowel Diseases.” N Engl J Med 383(27): 2652–2664.

Childs, C. J., H. M. Poling, K. Chen, Y.-H. Tsai, A. Wu, C. W. Sweet, A. Vallie, M. K. Eiken, S. Huang, R. Schreiner, Z. Xiao, A. S. Conchola, M. F. Anderman, E. M. Holloway, A. Singh, R. Giger, M. M. Mahe, K. D. Walton, C. Loebel, M. A. Helmrath, S. Rafii and J. R. Spence (2024). “Coordinated differentiation of human intestinal organoids with functional enteric neurons and vasculature.” bioRxiv: 2023.2011.2006.565830.

Clevers, H. (2016). “Modeling Development and Disease with Organoids.” Cell 165(7): 1586–1597.

Dong, Y., B. A. Johnson, L. Ruan, M. Zeineldin, T. Bi, A. Z. Liu, S. Raychaudhuri, I. Chiu, J. Zhu, B. Smith, N. Zhao, P. Searson, S. Watanabe, M. Donowitz, T. C. Larman and R. Li (2024). “Disruption of epithelium integrity by inflammation-associated fibroblasts through prostaglandin signaling.” Sci Adv 10(14): eadj7666.

Fatehullah, A., S. H. Tan and N. Barker (2016). “Organoids as an in vitro model of human development and disease.” Nat Cell Biol 18(3): 246–254.

Finkbeiner, S. R., X.-L. Zeng, B. Utama, R. L. Atmar, N. F. Shroyer, M. K. Estes and T. S. Dermody (2012). “Stem Cell-Derived Human Intestinal Organoids as an Infection Model for Rotaviruses.” mBio 3(4): e00159–00112.

Groschwitz, K. R. and S. P. Hogan (2009). “Intestinal barrier function: molecular regulation and disease pathogenesis.” J Allergy Clin Immunol 124(1): 3–20; quiz 21-22.

Guo, S., R. Al-Sadi, H. M. Said and T. Y. Ma (2013). “Lipopolysaccharide causes an increase in intestinal tight junction permeability in vitro and in vivo by inducing enterocyte membrane expression and localization of TLR-4 and CD14.” Am J Pathol 182(2): 375–387.

Hill, D. R. and J. R. Spence (2017). “Gastrointestinal Organoids: Understanding the Molecular Basis of the Host-Microbe Interface.” Cell Mol Gastroenterol Hepatol 3(2): 138–149.

Huch, M., J. A. Knoblich, M. P. Lutolf and A. Martinez-Arias (2017). “The hope and the hype of organoid research.” Development 144(6): 938–941.

Huizinga, J. D., C. M. McKay and E. J. White (2006). “The many facets of intestinal peristalsis.” Am J Physiol Gastrointest Liver Physiol 290(6): G1347-1349; author reply G1348-1349.

Kim, J., B. K. Koo and J. A. Knoblich (2020). “Human organoids: model systems for human biology and medicine.” Nat Rev Mol Cell Biol 21(10): 571–584.

Lancaster, M. A. and J. A. Knoblich (2014). “Organogenesis in a dish: Modeling development and disease using organoid technologies.” Science 345(6194): 1247125.

Liu, T., X. Li, H. Li, J. Qin, H. Xu, J. Wen, Y. He and C. Zhang (2024). “Intestinal organoid modeling: bridging the gap from experimental model to clinical translation.” Front Oncol 14: 1334631.

Ohland, C. L. and C. Jobin (2015). “Microbial activities and intestinal homeostasis: A delicate balance between health and disease.” Cell Mol Gastroenterol Hepatol 1(1): 28–40.

Sato, T., D. E. Stange, M. Ferrante, R. G. J. Vries, J. H. van Es, S. van den Brink, W. J. van Houdt, A. Pronk, J. van Gorp, P. D. Siersema and H. Clevers (2011). “Long-term Expansion of Epithelial Organoids From Human Colon, Adenoma, Adenocarcinoma, and Barrett’s Epithelium.” Gastroenterology 141(5): 1762–1772.

Sato, T., R. G. Vries, H. J. Snippert, M. van de Wetering, N. Barker, D. E. Stange, J. H. van Es, A. Abo, P. Kujala, P. J. Peters and H. Clevers (2009). “Single Lgr5 stem cells build crypt-villus structures in vitro without a mesenchymal niche.” Nature 459(7244): 262–265.

Schutgens, F. and H. Clevers (2020). “Human Organoids: Tools for Understanding Biology and Treating Diseases.” Annu Rev Pathol 15: 211–234.

Sonnier, D. I., S. R. Bailey, R. M. Schuster, M. M. Gangidine, A. B. Lentsch and T. A. Pritts (2012). “Proinflammatory chemokines in the intestinal lumen contribute to intestinal dysfunction during endotoxemia.” Shock 37(1): 63–69.

Spence, J. R., C. N. Mayhew, S. A. Rankin, M. F. Kuhar, J. E. Vallance, K. Tolle, E. E. Hoskins, V. V. Kalinichenko, S. I. Wells, A. M. Zorn, N. F. Shroyer and J. M. Wells (2010). “Directed differentiation of human pluripotent stem cells into intestinal tissue in vitro.” Nature 470(7332): 105–109.

Spence, J. R., C. N. Mayhew, S. A. Rankin, M. F. Kuhar, J. E. Vallance, K. Tolle, E. E. Hoskins, V. V. Kalinichenko, S. I. Wells, A. M. Zorn, N. F. Shroyer and J. M. Wells (2011). “Directed differentiation of human pluripotent stem cells into intestinal tissue in vitro.” Nature 470(7332): 105–109.

Spencer, N. J. and H. Hu (2020). “Enteric nervous system: sensory transduction, neural circuits and gastrointestinal motility.” Nat Rev Gastroenterol Hepatol 17(6): 338–351.

Uhlig, H. H. and F. Powrie (2018). “Translating Immunology into Therapeutic Concepts for Inflammatory Bowel Disease.” Annu Rev Immunol 36: 755–781.

van de Wetering, M., H. E. Francies, J. M. Francis, G. Bounova, F. Iorio, A. Pronk, W. van Houdt, J. van Gorp, A. Taylor-Weiner, L. Kester, A. McLaren-Douglas, J. Blokker, S. Jaksani, S. Bartfeld, R. Volckman, P. van Sluis, V. S. Li, S. Seepo, C. Sekhar Pedamallu, K. Cibulskis, S. L. Carter, A. McKenna, M. S. Lawrence, L. Lichtenstein, C. Stewart, J. Koster, R. Versteeg, A. van Oudenaarden, J. Saez-Rodriguez, R. G. Vries, G. Getz, L. Wessels, M. R. Stratton, U. McDermott, M. Meyerson, M. J. Garnett and H. Clevers (2015). “Prospective derivation of a living organoid biobank of colorectal cancer patients.” Cell 161(4): 933–945.

Vlachogiannis, G., S. Hedayat, A. Vatsiou, Y. Jamin, J. Fernández-Mateos, K. Khan, A. Lampis, K. Eason, I. Huntingford, R. Burke, M. Rata, D. M. Koh, N. Tunariu, D. Collins, S. Hulkki-Wilson, C. Ragulan, I. Spiteri, S. Y. Moorcraft, I. Chau, S. Rao, D. Watkins, N. Fotiadis, M. Bali, M. Darvish-Damavandi, H. Lote, Z. Eltahir, E. C. Smyth, R. Begum, P. A. Clarke, J. C. Hahne, M. Dowsett, J. de Bono, P. Workman, A. Sadanandam, M. Fassan, O. J. Sansom, S. Eccles, N. Starling, C. Braconi, A. Sottoriva, S. P. Robinson, D. Cunningham and N. Valeri (2018). “Patient-derived organoids model treatment response of metastatic gastrointestinal cancers.” Science 359(6378): 920–926.

Watson, C. L., M. M. Mahe, J. Múnera, J. C. Howell, N. Sundaram, H. M. Poling, J. I. Schweitzer, J. E. Vallance, C. N. Mayhew, Y. Sun, G. Grabowski, S. R. Finkbeiner, J. R. Spence, N. F. Shroyer, J. M. Wells and M. A. Helmrath (2014). “An in vivo model of human small intestine using pluripotent stem cells.” Nature Medicine 20(11): 1310–1314.

Workman, M. J., M. M. Mahe, S. Trisno, H. M. Poling, C. L. Watson, N. Sundaram, C. F. Chang, J. Schiesser, P. Aubert, E. G. Stanley, A. G. Elefanty, Y. Miyaoka, M. A. Mandegar, B. R. Conklin, M. Neunlist, S. A. Brugmann, M. A. Helmrath and J. M. Wells (2017). “Engineered human pluripotent-stem-cell-derived intestinal tissues with a functional enteric nervous system.” Nat Med 23(1): 49–59.

Yagi, S., A. Takaki, T. Hori and K. Sugimachi (2002). “Enteric lipopolysaccharide raises plasma IL-6 levels in the hepatoportal vein during non-inflammatory stress in the rat.” Fukuoka Igaku Zasshi 93(3): 38–51.

Ye, Q., J. Zhou, Q. He, R. T. Li, G. Yang, Y. Zhang, S. J. Wu, Q. Chen, J. H. Shi, R. R. Zhang, H. M. Zhu, H. Y. Qiu, T. Zhang, Y. Q. Deng, X. F. Li, J. F. Liu, P. Xu, X. Yang and C. F. Qin (2021). “SARS-CoV-2 infection in the mouse olfactory system.” Cell Discov 7(1): 49.

Yui, S., T. Nakamura, T. Sato, Y. Nemoto, T. Mizutani, X. Zheng, S. Ichinose, T. Nagaishi, R. Okamoto, K. Tsuchiya, H. Clevers and M. Watanabe (2012). “Functional engraftment of colon epithelium expanded in vitro from a single adult Lgr5+ stem cell.” Nat Med 18(4): 618–623.

