## Supplemental Table S1&2 for "Centimeter-scale, physiologically relevant intestinal organoids generated entirely from pluripotent stem cells"

Supplementary Materials

Table S1: Antibody information.

| NEON® 9-color DendronFluor® TSA kit (50~150 slides) | NEFP9100 |
| --- | --- |
| Rabbit Anti-Ki67 | CST 9027 |
| Rabbit Anti-Chromogranin A | Abcam ab283265 |
| Rabbit Anti-Muc2 | Abcam ab272692 |
| Rabbit Anti-CD31 | Abcam ab182981 |
| Rabbit Anti-a-SMA | CST 19245 |
| Rabbit Anti-Perilipin | CST 9349 |
| Rabbit Anti-CD68 | Abcam ab213363 |
| Rabbit Anti-E-cadherin | CST 3195 |

Table S2: qPCR primer information.

| IL6 F | AGACAGCCACTCACCTCTTCAG |
| --- | --- |
| IL6 R | TTCTGCCAGTGCCTCTTTGCTG |
| CCL2 F | AGAATCACCAGCAGCAAGTGTCC |
| CCL2 R | TCCTGAACCCACTTCTGCTTGG |
